# Evolution of Vesicle Shape Parameters Based on A Deep-Learning Based Image Analysis Framework Over Extended Time Periods

**DOI:** 10.64898/2026.11.03.764022

**Authors:** Tanmay Pandey, Rajni Kudawla, Sourav Negi, Varshan B S Jeyanth, Chetana Devarakonda, Deepika Singh, Aritra Chatterjee, Tripta Bhatia

## Abstract

A precise analysis of shape instabilities in cell-sized membrane compartments requires thorough quantification of the shape dynamics of individual components over extended periods. Herein, we have used phase-contrast microscopy to image giant unilamellar membrane vesicles (GUVs), which are widely used as model systems for cellular structures. GUVs were prepared in the fluid lamellar phase of 1-palmitoyl-2-oleoyl-sn-glycero-3-phosphocholine with and without cholesterol dissolved within a potato starch solution. The morphological changes and budding dynamics of individual GUVs with different chemical compositions were quantified using a deep-learning based image analysis framework over extended time periods, by using phase-contrast micrographs as input data. This automated framework integrates contour-based detection of the mother vesicle with precise segmentation of daughter compartments, thereby enabling explicit identification and tracking of mother and daughter vesicles at high temporal resolution. These findings further facilitate the measurement of the temporal evolution of vesicle morphometrics including area, and local membrane curvature during budding, that can be used as an input to develop computational models aimed at elucidating curvature-driven membrane remodeling in complex soft materials.

## 1 Introduction

Cellular morphometrics depend on complex membrane behaviors regulated by active biochemical processes, including protein activity and cytoskeletal rearrangements ^1–6^. Quantitative, temporal tracking, and contact-point curvature analysis over extended time periods is essential for understanding dynamic changes in cell morphology ^7,8^. However, analysis of the long temporal data for the label-free time-lapse images of evolving mother–daughter vesicle structures is not trivial.

The primary work presented here is an applicable imageanalysis framework for extracting physical parameters such as mean curvature and relaxation time scales. We have integrated the existing segmentation models into a physically constrained image-analysis workflow that enables quantitative measurements of membrane remodeling. The novelty of our approach does not lie in the development of a new deep learning architecture. Rather, the existing segmentation models are integrated into a physically constrained image-analysis workflow that enables quantitative measurements of membrane remodeling.

In this paper, we have combined contour-based detection of the vesicles’ contour with segmentation of daughter compartments, followed by geometry-based filtering, temporal tracking, and contact-point curvature analysis over extended time periods. We have presented the dataset capturing the complete spatio-temporal evolution of shape changes in vesicles composed of the lipids 1-palmitoyl-2-oleoyl-sn-glycero-3-phosphocholine (POPC) and POPC-Cholesterol. Our approach involves using a reference membrane compartment such as giant unilamellar vesicles (GUVs) of POPC and POPC-Cholesterol prepared in the potato starch solution at a concentration of 0.5 mg/ml. The inner and outer solutions are composed of the same solution (with the same refractive index), and light passing through the interior solution does not experience any phase shift relative to the exterior solution. In contrast, the high refractive-index contrast between the membrane and the surrounding medium makes the membrane appear as a dark boundary, as shown in Supplementary Figure 1, only at the equatorial plane of the GUV membrane, which remains sharply in focus. This allows estimating the overall shape and size of the entire GUV by measuring the equatorial area of a spherically symmetric vesicle. GUVs generally have diameters of 10 − 100 *µ*m, which permits direct visualization with phase-contrast microscopy (PCM) ^9^. These vesicles consist of a single lipid bilayer and are typically formed by electroformation or gelassisted swelling. GUVs allow for systematic study of membrane shape changes—such as budding and tubulation—resulting from a balance between membrane elasticity, surface tension, lipid interactions, and the surrounding solution. External chemical or light-based perturbations can further induce shape changes by altering membrane tension and curvature, as shown in numerous GUV studies. Their adjustable membrane composition enables exploration of how membrane physical properties respond to external physicochemical signals. ^8,10–12^

The manuscript uses deep-learning-based segmentation tools. Accurate estimation of the morphological dynamics of GUVs require meticulous detection of individual vesicle components at high temporal resolution over extended time durations. Automated image analysis frameworks allow for precise quantification of individual frames within large data sets and therefore are highly useful for such measurements. In recent years, several software and programming tools have been developed to automate quantitative analysis of microscopy data for living specimens, though most of them require fluorescence microscopy rather than phase-contrast imaging as an input ^13,14^. Additionally, these frameworks focus on population-level descriptors rather than individual-vesicle shape dynamics over time. GUV-AP, a Fiji-based macro, uses circular Hough detection to rapidly locate vesicles and quantify membrane dye incorporation and protein-binding intensity across large fields of view, but assumes near-circular geometry and is not designed for tracking internal compartments ^14^. DisGUVery, an open source python-based framework for quantitative analyses of GUVs, combines multiple detection algorithms, including circular Hough and contour-based approaches with modules for membrane fluorescence, shape, and internal-content analysis, and can explicitly support non-spherical vesicle detection via a dedicated membrane-segmentation algorithm ^13^. However this framework, does not address the simultaneous identification and temporal linking of nested mother–daughter compartments ^13^. Deep-learning approaches have also been applied to quantitative GUV image analysis — for instance, convolutional neural networks have been used to automatically classify vesicle morphology and membrane-binding state in confocal fluorescence data; but these efforts similarly target single-compartment classification rather than multi-object segmentation and tracking within a single mother vesicle..

In this study, we present an automated framework for image analysis that integrates shape-based detection with advanced deep learning techniques for image segmentation and tracking. The framework presented here is built for label-free phasecontrast time-lapse data, and combines contour-based mother-vesicle detection with deep-learning instance segmentation (Cellpose/StarDist ^15,16^) of daughter compartments and enables simultaneous tracking of both the outer boundary and the internal budding population at high temporal resolution. These methodologies establish automated, scalable workflows that facilitate robust, time-resolved quantification of membrane-compartment shape dynamics. Application of this proposed framework facilitates the accurate detection of vesicle shape descriptors at high spatiotemporal resolution and can be used for comparative analysis of membrane remodeling for GUVs with different concentrations that can be broadly applied to various types of image data and are not necessarily limited to phase-contrast microscopy

## 2 Materials and Methods

### 2.1 Materials

1-palmitoyl-2-oleoyl-sn-glycero-3-phosphocholine (POPC) lipid (Catalog number 850457P), 1,2-Dioleoyl-sn-glycero-3-phosphocholine (DOPC) lipid (Catalog number 850375P), Cholesterol (Catalog number C8503) and 1,2-dihexadecanoylsn-glycero-3-phosphocholine (DPPC) lipid (Catalog number 850355P) were purchased from Avanti Polar Lipids in powder form and dissolved in chloroform to prepare a stock solution at a concentration of 1 mg/ml. The lipid stock solutions are kept at −20°C. Alpha-amylase (AA) from Malt was purchased from HI media (Catalog number 70 GRM638), Starch soluble GR (CAS No. 9005-84-9) and for confocal experiments nile red dye (Catalog number 7385-67-3) and Fluorescein isothiocyanate (FITC) dye (Catalog number 27072-45-3) were obtained from Sigma-Aldrich. To prepare the starch stock solution in Milli-Q water, 5 mg of soluble starch powder was weighed and added to 7-8 ml of warm Milli-Q water. The mixture was then heated to 60°C - 70°C with a magnetic stirrer rotating at 500 rpm until the starch dissolved. Subsequently, the volume was adjusted to 10 ml with Milli-Q water in a Falcon tube. The tube was placed on a rotator for 5 to 10 minutes to ensure uniform mixing. To probe the biochemical activity of AA hydrolysis in the bulk starch solution, 3,5-dinitrosalicylic acid (DNS) analysis was performed to quantify reducing sugar production in the external medium. ^17,18^

### 2.2 Methods

GUVs composed of (i) 1-palmitoyl-2-oleoyl-sn-glycero-3-phosphocholine (POPC), (ii) POPC:Cholesterol (9:1) and (iii) DOPC:DPPC:Cholesterol (0.35:0.35:0.30) were prepared using the electroformation method described by Angelova and Dimitrov ^19^. These GUVs were prepared in a 0.5 mg/ml potato starch solution at 37°C. Sample cells for GUVs experiments are observed under a phase-contrast microscope (Zeiss Axio Observer 5) equipped with a 20× dry objective (0.4 N.A.) and a Zeiss Axiocam 506 mono camera. All experiments in the present study were conducted at room temperature 24 °C. The phasecontrast microscope allows the observation of cellular structures and components without the need for fluorescence. Since its development in the early 1930s, phase-contrast microscopy represented a major breakthrough. ^9^ ImageJ is used for manual tracking. In HPLC experiments, Refractive Index Detector (RID) chromatogram, mV is the detector’s output signal (Y-axis), which is a voltage proportional to the difference in refractive index between the sample eluting from the column and the pure mobile phase in the reference cell. A peak’s height in mV reflects analyte concentration, and its area (mV·s) is used for quantification. The X-axis is retention time.

### 2.3 Experimental data

Figure 1 presents time-lapse phase-contrast images illustrating the morphological changes of starch-loaded POPC GUVs following exposure to 2.5 *µM* AA. The experiment initiates at *t* = 0, marking the point at which the enzyme is introduced to the exterior compartment of the GUVs. The left panel exhibits the raw micrographs of the vesicles, while the right panel depicts the detected contours of both the mother vesicles (outermost) and daughter vesicles (inter luminal) within each time-series dataset. Time stamps are included with the images. Each temporal frame was subjected to analysis for the concurrent detection of mother vesicle contours and the identification of contained daughter vesicles. It was observed that within 30 minutes of adding AA, the vesicles began to transform in shape, a process that persisted up to a few hours. This evolution resulted in the mother vesicle becoming increasingly crowded with internal quasi-spherical buds, as illustrated in Figure 1. For vesicles or buds of spherical shape, the relationship between their equatorial area (*A*) and the total surface area (*S*) is expressed as *S* = 4*A*. The left panel in Figure 1 highlights the raw image, and the right panel shows the equatorial areas defined by the boundary of the mother vesicle and those of the individual daughter vesicles contained within it, with the cyan color. To explore the robustness of our observations, we performed experiments using one-component POPC vesicles, a binary lipid system composed of POPC: Cholesterol (9:1), and a ternary lipid system composed of DOPC:DPPC:cholesterol (0.35:0.35:0.30), forming GUVs under similar conditions. Upon addition of AA at the same concentration, we observed inward budding and membrane remodelling. The membrane domains are observed for DOPC:DPPC:cholesterol (0.35:0.35:0.30) GUVs. As shown in Figure 2, inward buds are observed for the three lipid compositions, including the more complex membrane systems, such multicomponent membranes displaying for instance phase separation. The domains are present in the internal buds/daughter compartments and shown by an arrow in Figure 2.

**Fig. 1.**
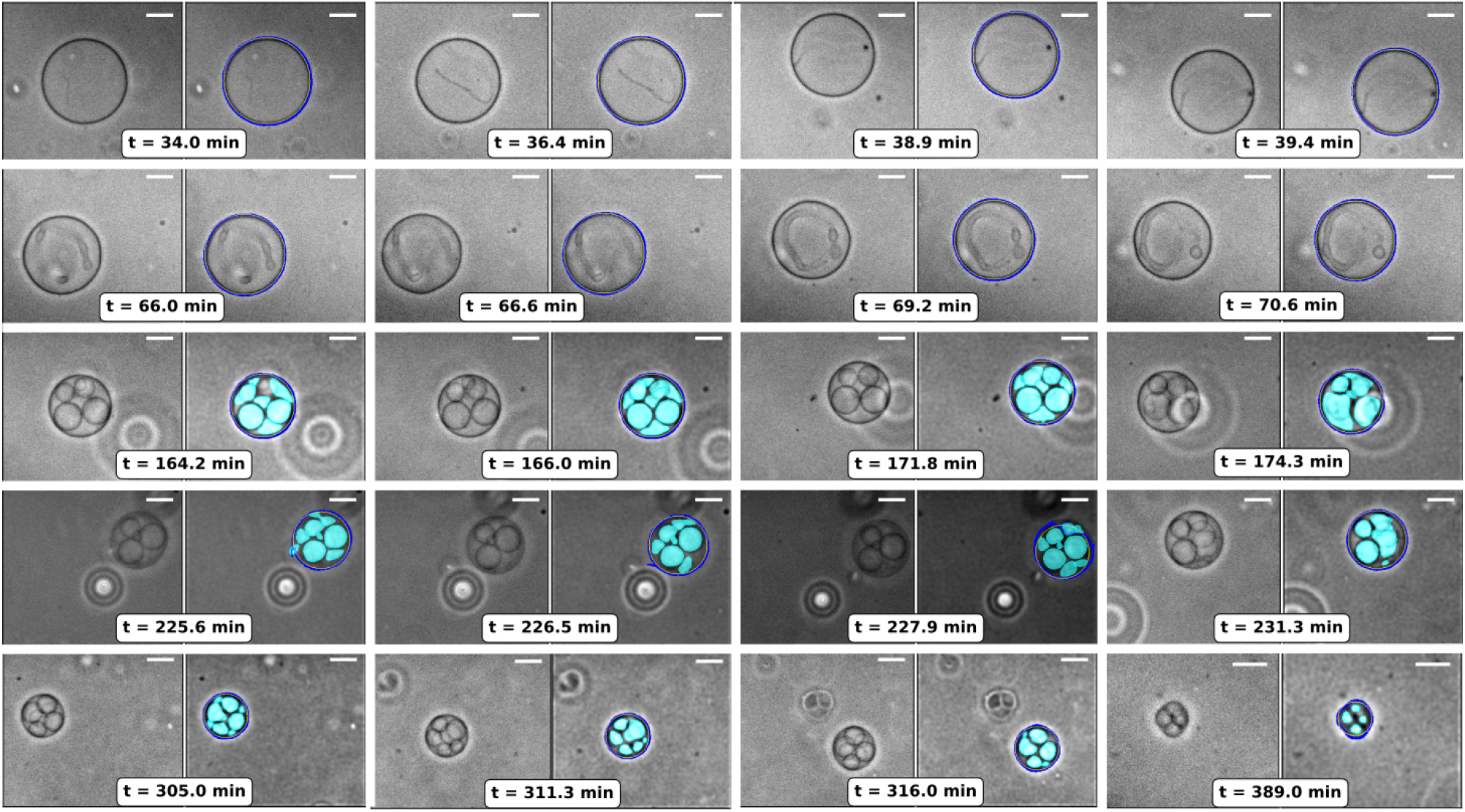
The time-series montage shows morphological changes in a vesicle over time. On the left, raw images with equatorial regions of interest are displayed, while on the right, the detected mother vesicle contours (dark black) and bud segmentation masks (cyan) are shown side by side. Scale bars represent 10 *µm*.

**Fig. 2.**
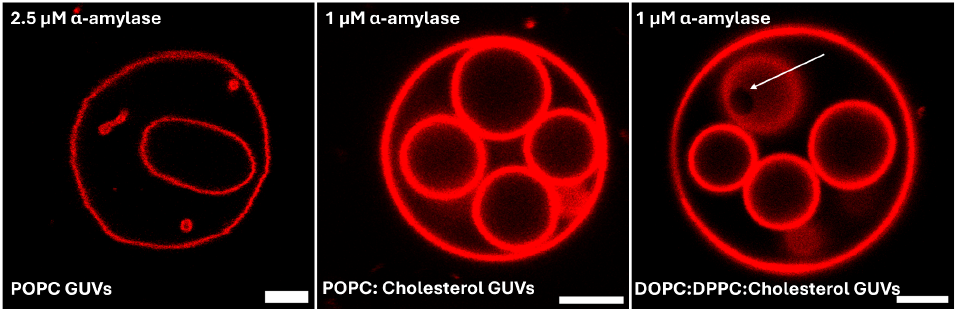
Three different lipid mixtures and the fluorescence images of GUVs showing membrane budding and phase separation under enzymedriven osmotic deflation. The DOPC:DPPC:cholesterol GUV display dark region lacking the fluorescence dye shown by an arrow, characteristic of lateral lipid phase separation. GUVs exhibit prominent inward budding/internal compartmentalisation, characterized by discrete, approximately circular membrane structures located within the parent GUV. The phase-separated domains remain confined to and continuous with the parent membrane and therefore differ morphologically from enclosed internal buds. The first image is taken after 1 hour 3 minutes, second image is taken after 3 hours 10 minutes and third image is taken after 3 hours 30 minutes, after adding the enzyme AA. The scale bar is 5 *µ*m and applied to all the images. Membrane dye is Nile Red.

The manuscript analyzes neck curvature between mother and daughter compartments and between daughter compartments, which implies connectivity through membrane necks. Figure 2 shows images of internal vesicles inside the mother GUV. To determine whether the intra-luminal daughter vesicles formed during membrane remodelling remained connected to the mother vesicle, a series of fluorescence-labelling experiments were performed. GUVs were prepared in 0.5 mg/mL starch solution, and AA was added externally shown in Figure 2. Nile Red was added before confocal imaging. Confocal z-stack imaging was performed at 24 °C to visualise both membrane organisation and the formation of internal buds/compartments. The intra-luminal vesicles are connected to the mother vesicles via the membrane neck shown in Figure 2; otherwise, the membrane dye will not reach the internal compartments when added in the bulk solution. The introduction of the enzyme AA induced a gradual contraction of the mother vesicles over time, an increase in crowding internally, and the formation of multiple internal compartments, shown in Figure 1.

### 2.4 Osmotic deflation

Figure 3 shows HPLC-RID chromatogram of 500 ppm starch hydrolysed by 2.5 µM AA at 25 °C. The identified products in the bulk solution are glucose, maltose, maltotriose, maltotetraose, maltopentaose, maltohexaose, and maltoheptaose. Isocratic gradient conditions having mobile phase of acetonitrile and water in ratio of 70:30 (v/v) were used at a flow rate of 0.8 mL/min. Aminopropyl (NH2) column having dimensions of 250 x 4.6 mm having pore size of 5 *µ*m used having run time of 40 min. Column temperature was maintained at 30 °C whereas refractive index detector was maintained at 35 °C. The Figure 4 (top image) shows an initial rapid increase in the overall reducing sugar concentration within the first 30 minutes. This is attributed to the introduction of the enzyme AA in the exterior of the GUVs and AA-catalyzed hydrolysis of potato starch during a 300 minutes incubation. Figure 4 displays Phase-Contrast microscopy images of POPC GUVs acquired at 34.5, 129, and 263 minutes, respectively, of starch hydrolysis by AA. The membrane necks are shown in phase-contrast images in Figure 4 where LS, SS denote the neck between the large-small vesicles and small-small vesicles, respectively. The greater concentration of reducing sugar in the external bulk solution relative to inside the GUVs leads to osmotic deflation and the formation of inbuds.

**Fig. 3.**
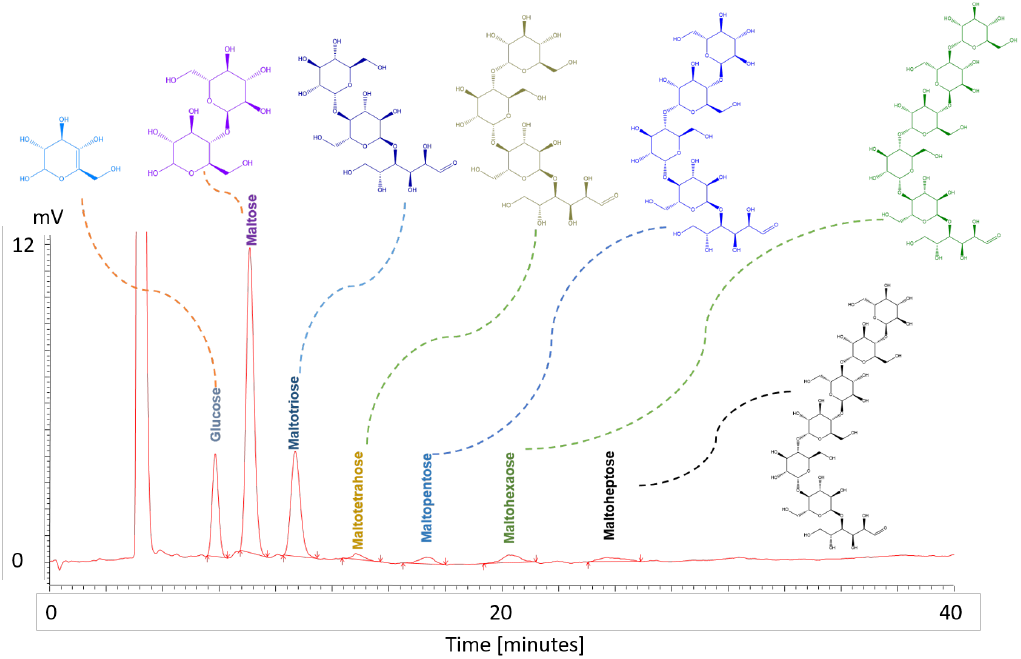
HPLC–RID chromatogram of 500 ppm starch hydrolyzed by 2.5 *µ*M AA at 25 °C. The identified products are glucose, maltose, maltotriose, maltotetraose, maltopentaose, maltohexaose, and maltoheptaose.

**Fig. 4.**
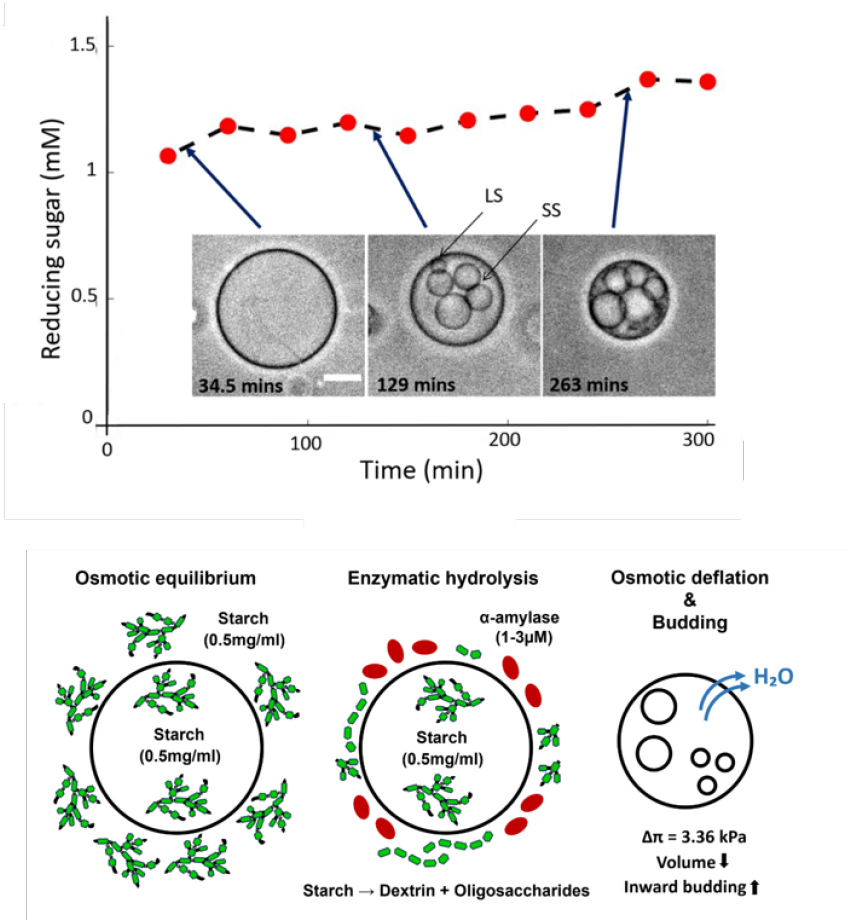
Image shows a time-dependent increase in reducing sugar concentration observed during AA-mediated starch hydrolysis at a concentration of 2.5 *µ*M AA. Phase-contrast microscopy images of the same POPC GUV are taken at 34.5, 129, and 263 minutes during starch hydrolysis in the bulk solution outside the GUVs. The scale bar is 10 *µm* and applies to all images. LS and SS show the necks between the large-small and smallsmall vesicles, respectively. The schematic shows osmotic imbalance and inward budding in GUVs.

The schematic shows that the formation of soluble sugars during enzymatic hydrolysis increases the osmolarity of the external solution surrounding the GUVs. This generates an osmotic pressure difference between the vesicle interior and the external medium. The osmotic pressure difference (ΔΠ) generated across the vesicle membrane can be estimated using the van’t Hoff relation with a difference in solute concentration across the membrane (Δ*C*).

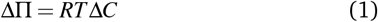

where *R* is the universal gas constant having value 8.314 Jmol^−1^ K^−1^, *T* is the absolute temperature (298 K). Using the experimentally determined concentration of reducing sugars is varying between 1 − 1.5 mM as shown in the Figure 4 during the reaction, the osmotic pressure difference is varying between 2.6 − 3.4 kPa, respectively, from 30 − 300 minutes.

The introduction of the enzyme AA induced a gradual contraction of the mother vesicles over time, an increased internal crowding, and the formation of multiple internal compartments. Thus, the shape changes can be attributed to the accumulation of reducing sugars outside the GUVs, as found in the HPLC data shown in Figure 3. Previous studies have validated that shape instabilities in GUVs may arise from membrane structural or compositional asymmetries, inner-outer leaflets area mismatches, or other non-equilibrium processes ^10–12^. The work reported in this paper is about the image-analysis framework to extract the membrane curvatures and membrane surface area of the intraluminal daughter vesicles formed during membrane remodeling by different sugars in contact with the outer leaflet of the GUV membrane. We have integrated the existing segmentation models into a physically constrained image-analysis workflow that enables quantitative measurements of membrane remodeling.

## 3 Automated Image Analysis Framework for Vesicle Shape Analysis

In this section, we have described the different stages of the analysis, including preprocessing, mother-vesicle detection, daughtervesicle segmentation, geometry-based filtering, temporal tracking, and curvature extraction..

### 3.1 Vesicle Analysis Initialization, Figure 5 - Phase 1

Time-series images of the GUVs were obtained through phase-contrast microscopy and assembled into stacks, with each image representing a specific time point. Any delay in data acquisition was taken into account while syncing the individual frames in the corresponding stream of images.

**Fig. 5.**
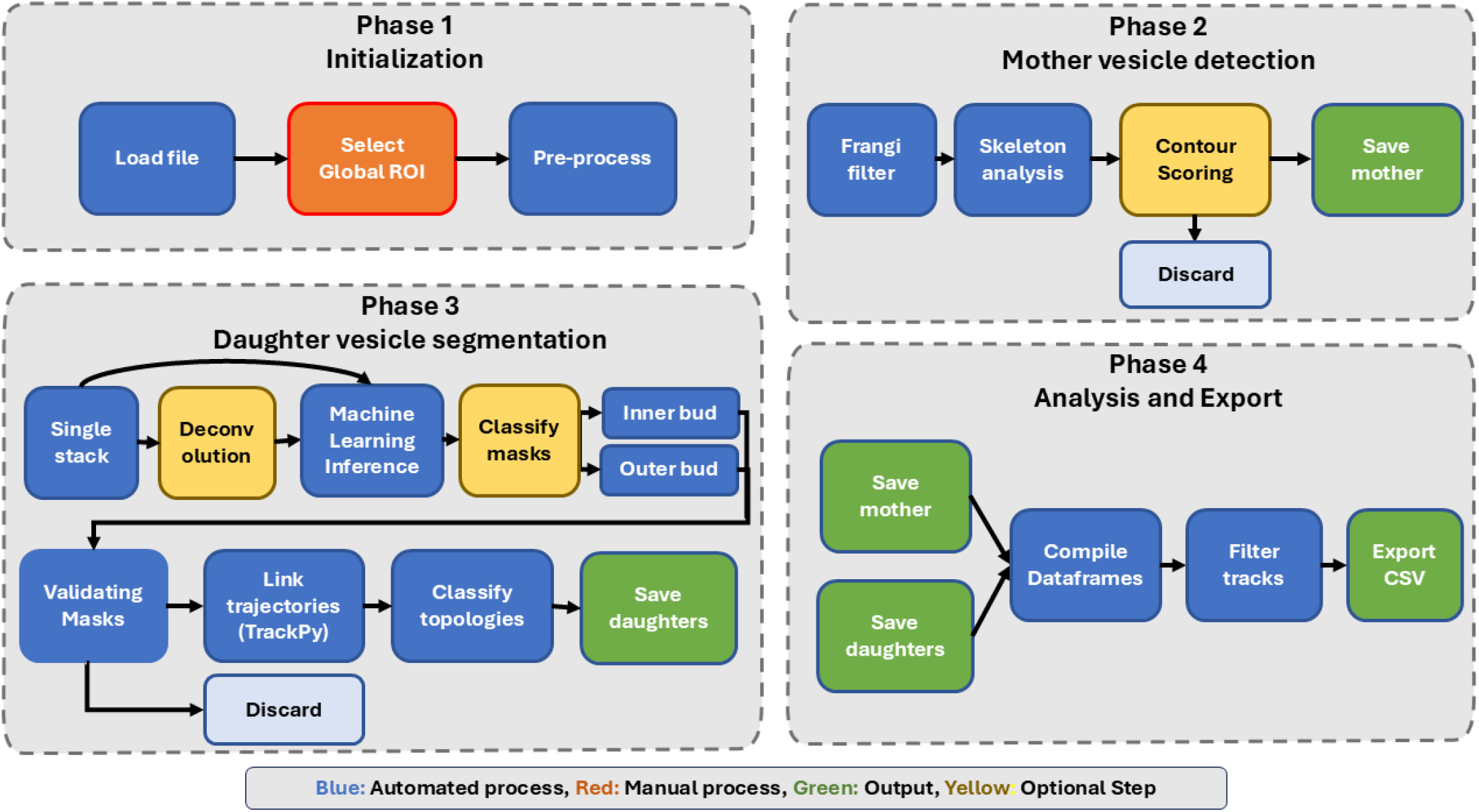
Schematic showing the image analysis workflow employed in this study. The workflow includes four phases: initialization and pre-processing, detection of mother vesicles, segmentation and tracking of daughter vesicles, and data analysis with export. Automated steps are marked in black, manual steps in red, optional steps in yellow, and saved outputs are indicated in green.

#### Preprocessing of Acquired Images

We imported the timelapse-based image stacks into a Python environment (version 3.10) and then converted to NumPy arrays. Each stack was interactively cropped to a specific region of interest (ROI) containing a GUV, thereby excluding the background regions and any neigh-boring vesicles that could interfere with segmentation and tracking (Figure 5 - Phase 1), which also helped in array size reduction. ROI selection was performed uniformly across all frames; however, when required, frame-dependent boundaries were also used to account for vesicle drift during prolonged time-lapse data acquisition.

To enhance segmentation robustness, additional preprocessing steps were employed prior to the vesicle detections. Intensity fluctuations caused by uneven illumination and shifts were minimized by normalizing the intensities across all frames. In some datasets, additional contrast enhancements and background suppression were performed using an interactive GUI.

### 3.2 Mother Vesicle Detection, Figure 5 - Phase 2

#### Automated Detection of Mother Vesicle Contour

We segmented the mother vesicle using a multi-stage, contour-based strategy that analyzed each frame independently while using temporal information from preceding frames to preserve continuity of vesicle shape and position (Figure 5 - Phase 2). We first smoothed the images with a Gaussian filter to reduce noise ^22–24^, then applied a gradient operator to identify membrane edges, treating edges as gradient discontinuities ^25,26^. This step combined a firstderivative wavelet transform ^27^ with a Sobel operator ^28,29^ wherever applicable.

#### Dynamic Detection of Contours using Weighted Scoring

To ensure seamless detection, we assigned each candidate contour a weighted score based on three parameters: (i) circularity (*C* = (4*πA/P*^2^), where *P* is the perimeter, and *A* is the enclosed area of the detected contour), (ii) vesicle area, and (iii) temporal consistency. An ellipse was fit to each contour and a distance-based similarity metric was then employed to quantify its deviation from the detected contour, preventing fragmented detections. We applied this size-based score specifically to contours having a radius greater than 5 *µm*, thereby automatically excluding very small or fragmented contours that might otherwise be mistaken for artifacts. To maintain temporal consistency, we compared each candidate contour to the previously identified mother vesicle by tracking centroid displacement and changes in effective radius across frames, and penalised large shifts in position or size, since these do not match the gradual, continuous changes typical of the mother vesicle under our experimental conditions. The final contour for each frame was subsequently selected using a weighted combination of shape, size, and temporal scores.

### 3.3 Daughter Vesicle Segmentation, Figure 5 - Phase 3

#### Daughter Vesicle Contour Detection

The accuracy of mother- and daughter-vesicle detection depends strongly on image quality and contrast. For image stacks with reduced contrast or blurred boundaries, we deconvolved all frames uniformly using an iterative Richardson–Lucy approach ^30^, with a controlled iteration count and background regularisation, before passing them to the detection framework (Figure 5 - Phase 3). We chose the deconvolution parameters, specifically the kernel width and the Gaussian point spread function (PSF) — interactively via an interactive graphical user interface.

We segmented daughter vesicles using the deep-learning-based instance segmentation framework StarDist ^16,31^, which models region boundaries as star-convex polygons and is particularly effective at resolving closely spaced or weakly contrasted membrane-bound structures. StarDist was selected over alternative frameworks due to its favourable noise robustness and segmentation accuracy, as highlighted in Supplementary Figure 2a-d.

We normalised raw frames before segmentation to minimise the effect of background fluctuations and then used them to generate segmentation masks using a pretrained StarDist model. Only the masks lying inside the mother-vesicle contour were retained, and those lying outside this boundary were discarded. This geometry-informed filtering provides us with an automated basis for subsequent tracking and contact analysis of in-buds.

### 3.4 Analysis and Export, Figure 5 - Phase 4

#### Tracking and Filtering Masks

We performed temporal linking of the daughter vesicles using the TrackPy library ^33^, which implements nearest-neighbour particle tracking with memory to accommodate intermittent segmentation failures. We fixed the tracking reference frame to the mother vesicle to remove overall drift across the time series, and excluded trajectories shorter than five frames from further analysis. Any remaining artefacts were manually removed during postprocessing.

We quantified vesicle interactions by computing the minimum Euclidean distance between successive vesicle contours, using nearest-neighbour queries via KD-tree search as implemented in SciPy ^34^. We applied a contact threshold of 10 pixels to identify direct contacts between daughter vesicles and the mother membrane, and independently queried per-daughter KD-trees to identify contacts between neighbouring pairs of daughter vesicles. We selected 10 pixels as the contact threshold because the number of detected contacts approaches a plateau at approximately this threshold, with only a relatively small increase in the number of contacts for thresholds above 10 pixels (Supplementary Figure 2f). Additionally, this 10-pixel threshold helps exclude artefacts that would interfere with daughter-mother contact detection. At each contact site, we estimated the local membrane radius of curvature using the three-point contour geometry, fitting a circle through the contact point and its two nearest contour neighbours. Together, this analysis yields stable vesicle trajectories and quantitative geometric observables suitable for comparison across vesicle sizes and membrane compositions.

#### Normalization of area and time

To facilitate comparisons of budding dynamics across vesicles and experimental conditions, the mother-vesicle and bud areas, as well as the recording time, were normalized. The mother-vesicle area was normalized by its 95th percentile value to minimize the influence of unusually large area values, whereas the combined bud area was normalized by its maximum observed value. Time was normalized by the total duration of the budding event, defined as the maximum recorded time (*t*_max_). These normalization procedures account for differences in absolute vesicle size and recording duration, enabling comparisons of relative changes in vesicle morphology and the progression of budding across different vesicles and experimental conditions.

## 4 Results

### 4.1 Model Validation for robustness against noise

Next, a more quantitative assessment of segmentation uncertainty, error in daughter-vesicle detection, and differences in performance between the segmentation models used is discussed. This is especially important since the daughter-vesicle analysis appears to be the most challenging part of this work. We evaluated the efficiency of our segmentation-based framework by progressively applying increased levels of Gaussian noise to the images to assess both segmentation performance and the robustness of the subsequent vesicle quantification. Intersection over Union (IoU) metric was primarily chosen as an important criterion for model selection.

We tested multiple segmentation-based approaches for our study, including watershed and active-contour-based detection, along with existing frameworks like Cellpose and StarDist. Among all the tested approaches, StarDist showed the highest overlap with the reference annotations at low noise and retained measurable segmentation performance at intermediate noise levels (Figure 6a) and was therefore selected for the subsequent vesicle segmentation and analysis.

**Fig. 6.**
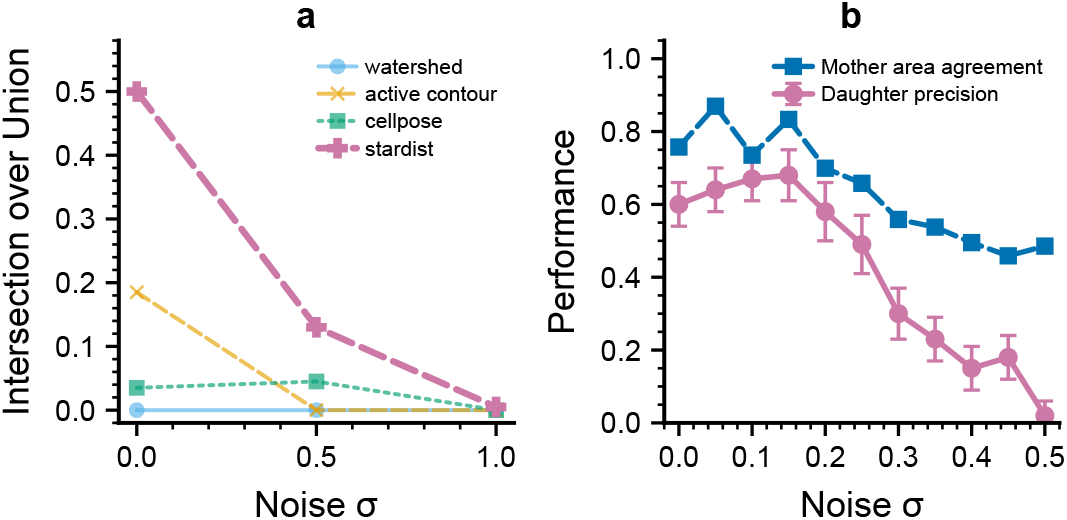
Validation of the segmentation pipeline by measuring segmentation metrics under increased image noise. (a) Variation in Intersection over Union (IoU) for the tested segmentation models at increasing noise levels. (b) Variation in Daughter vesicle precision and mother vesicle area agreement for the selected StarDist-based pipeline under increased noise levels.

Next, we evaluated how increasing image noise affected the quantitative measurements obtained from the selected pipeline, i.e StarDist. Daughter vesicle precision remained approximately 0.6– 0.7 at low noise levels and decreased progressively with increasing noise, with a pronounced reduction above *σ* ≈ 0.2 (Figure 6b). In parallel, we quantified the agreement between the segmented and reference mother-vesicle areas. This agreement remained relatively high at low noise levels and decreased progressively as the noise increased, indicating that the detection of small daughter vesicles is more sensitive to image degradation than the recovery of the larger mother-vesicle morphology.

Additional validation metrics provided in Supplementary Figure 2a-d further reinforce the choice of StarDist for our segmentation pipeline. The mother centroid error remained small (< 15%) at low noise levels but increased substantially for *σ >* 0.25. Additionally, we also compared the Hausdorff discrepancy and vesicle quantification deviation obtained using the different segmentation approaches (Supplementary Figure 2b,c). These measurements provide complementary assessments of contour localisation and the propagation of segmentation errors into down-stream vesicle quantification. Together, the IoU comparison and subsequent robustness analysis support the use of StarDist for vesicle segmentation. StarDist achieved the highest true-positive (TP) count among the tested approaches (Supplementary Figure 2d and Supplementary Table 1b), motivating its use for the subsequent daughter-vesicle analysis and characterising the range of image quality over which reliable morphological measurements can be obtained.

As a complementary, real-data check of pipeline reliability, we compared automated tracking results against independent manual annotation (ImageJ) for three representative POPC GUVs exposed to 2.5 µM alpha-amylase (V1–V3), quantifying agreement via Pearson’s correlation coefficient (*ρ*). As summarised in Supplementary Table 1a, we found strong positive correlations (*ρ >* 0.85) between the automated and manually tracked mother-vesicle area, with statistically significant *p*-values (*p <* 0.05) across all three vesicles. Although the distribution in measured daughter-bud area displayed greater variability, reflecting the added segmentation complexity of densely budded, highly dynamic regions, correlations remained strong (*ρ >* 0.7) for all three cases ^35^, and were particularly close for V1 and V3. Taken together with the noise-robustness analysis above, these results indicate that the automated framework not only tolerates synthetic image degradation but also reproduces independent manual measurements on real experimental data, supporting its suitability for high-throughput, long-term GUV shape analysis.

[Reviewer 3, major comment 9] The correlation with manual measurements alone is not sufficient to establish the accuracy of daughter-vesicle detection. We have now extended the framework validation to an ‘object-level analysis’ using manually annotated daughter vesicles as reference. For each validation frame, the predicted daughter objects were matched to corresponding manually annotated objects using an intersection-over-union (IoU) threshold of 0.5. Matched objects were classified as true positives (TP), while unmatched predicted objects and unmatched reference objects were classified as false positives (FP) and false negatives (FN), respectively. Precision, recall and F1 score were then calculated from these object-level detections and summarised in Supplementary Table 1b. We report TP, FP, FN, precision, recall and F1 score as object-level daughter-detection metrics for different segmentation algorithms as reported in Supplementary Table 1b and Supplementary Figure 2d. Together, these analyses provide quantitative measures of missed and spurious daughter detections and complement the previously reported temporal correlation analysis.

### 4.2 Analysis of Time-Lapse Vesicle Images

Figure 1 depicts the temporal evolution of vesicle morphology over a period of six hours along with the corresponding mother vesicle contours and segmented daughter vesicles detected using our framework. The left panel exhibits the raw micrographs of the vesicles, while the right panel demonstrates the detected contours of the mother and daughter vesicles within each time-series dataset. Collectively, the figure illustrates the reliability of our framework, which can precisely identify the mother vesicle contour throughout the time course of the experiment as well as detecting the individual daughter vesicles (buds) even within crowded conditions. We plotted the temporal evolution of the equatorial area (*A*) of the mother vesicle and daughter vesicles for the representative GUV depicted in Figure 1, resolved at high temporal resolution within a single small window (≈ 14 minutes) as shown in Supplementary Figure 3. For the mother vesicle, the equatorial area (*A*) is defined by the area of the region enclosed within the detected contour, while for the daughter vesicles we plotted the combined area covered by all such vesicles within a single image frame. These results provide evidence that the custom image segmentation framework can reliably monitor and quantify morphological changes in vesicles throughout the experiment across multiple independent recordings.

To test the efficacy of our custom-developed framework for robust segmentation and long-term tracking of GUV shape, we used experimental data that captured the entire shape evolution of these vesicles, from a single mother vesicle to the formation and growth of several in-buds (daughter vesicles). The observed increase in both the number and size of daughter vesicles over time validates the deep-learning–assisted segmentation framework’s capacity to accurately resolve overlapping and closely spaced structures, highlighting the temporal stability and geometric precision of the automated pipeline.

The complete temporal evolution of normalized *A* for the mother and daughter vesicles are shown in Figure 7 for 22 different GUVs varying over the concentration of alpha-amylase used in the outside solution and two different lipid types.

**Fig. 7.**
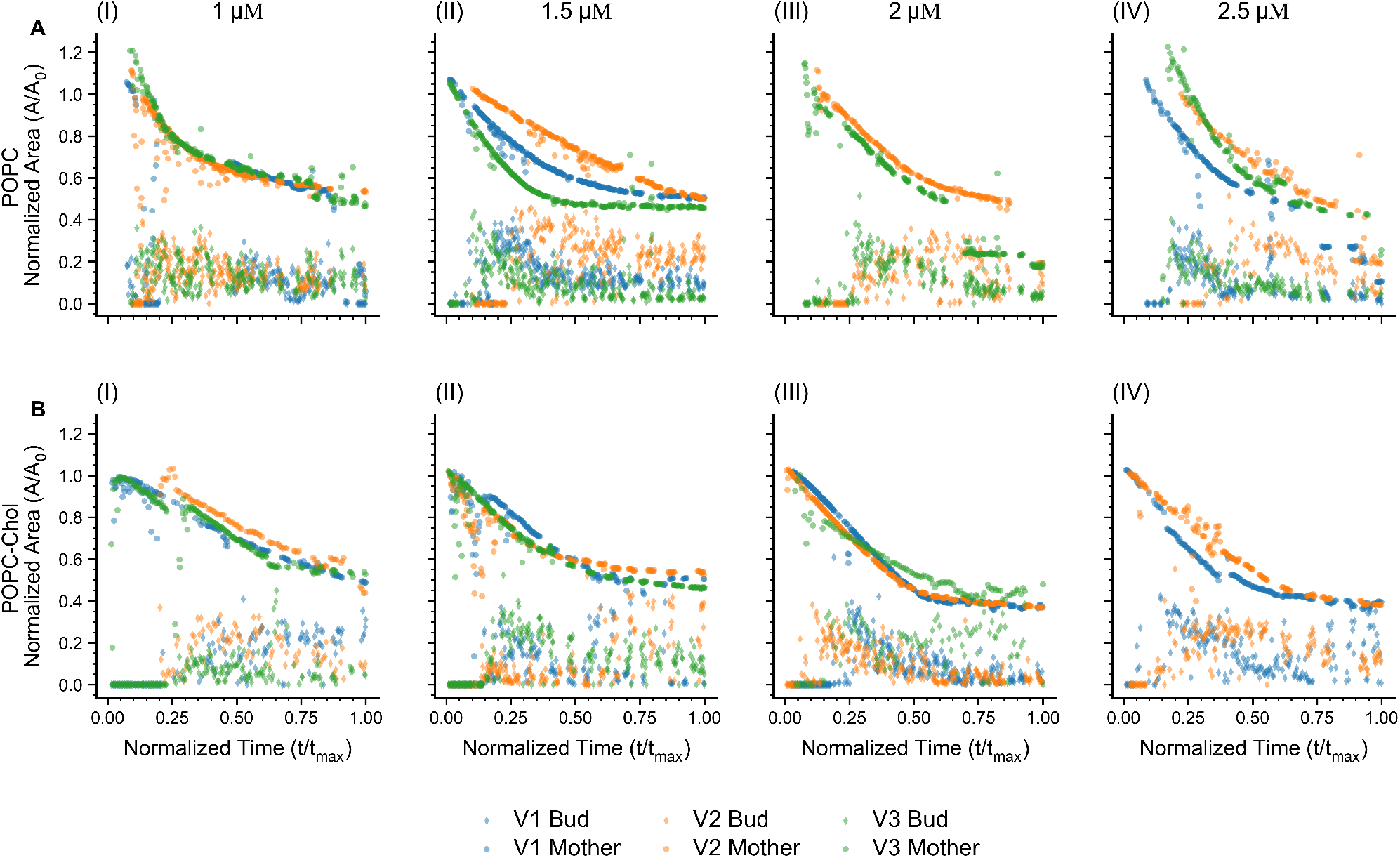
Evolution of normalized membrane area during vesicle budding. Normalized membrane area is plotted as a function of normalized time (*t/t*_max_) for representative (A) POPC and (B) POPC–Chol vesicles. Subpanels (I)–(IV) correspond to alpha-amylase concentrations of 1, 1.5, 2, and 2.5 *µ*M, respectively. Data from three representative vesicles (V1–V3) are shown, with circles representing the mother vesicle and diamonds representing the corresponding bud. Mother vesicle area was normalized by its 95*th* percentile area (*A*_0_), whereas bud area was normalized by its maximum observed area. Time was normalized by the total duration of the budding event (*t*_max_), enabling comparison of budding dynamics across vesicles and experimental conditions. A total of 22 vesicles were analyzed across all groups, with two compositions and four concentrations.

The (*A* = 0) in Figure 7 for a few specific frames, at the beginning of the experiments, are related to the absence of buds. Our automated framework can track these measurements (every 5^th^ frame) in the recorded data, reducing the size of the data to be processed. Our results highlight a time-dependent exponential decay in *A* for the mother vesicle, whereas the combined internal vesicle exhibits bounded exponential growth, asymptotically approaching a value close to the final mother vesicle area.

### 4.3 Extraction of Mean Curvature of the Membrane Neck

In order to quantify dynamic shape changes in the mother vesicle morphology during in-bud (daughter vesicle) formation over time, we computed specific curvature-based parameters from our analysis. For computing vesicle membrane curvatures, we select a convention where a positive sign is assigned to the outwardly generated curvature and a negative sign to the curvature of the inward-facing shapes. Supplementary Figure 4a displays a GUV with spherical symmetry. In contrast, the vesicles in Supplementary Figure 4b-e are non-spherical and exhibit inward-directed structures. Supplementary Figure 4f-j show, in green, the detected outermost contours of the mother vesicles corresponding to Supplementary Figure 4a–e, respectively. Supplementary Figure 4k–o present the normalized local curvature values obtained along the vesicle contours shown in Supplementary Figure 4a-e, respectively. As illustrated in Supplementary Figure 4f-i the contours of the mother vesicle can be accurately identified for various different morphologies at different stages of their shape evolution from the microscopy images. These contours were then used to quantify the spatial variations in local curvature along the detected boundary, as shown in Supplementary Figure 4k–o. Normalized curvature was defined using the relation *κ*_*n*_ = *κ/κ*_max_. The normalized local curvature [−1, 1] at each sample point of the contour is calculated, and its spatial variation along the vesicle boundary is depicted with the colorbar scale shown in Supplementary Figure 4. These results demonstrate the accuracy of contour detection and its applicability to various vesicle morphologies under static conditions. The comprehensive methodology for mapping the spatial variation of membrane curvature on the entire vesicle’s surface is discussed separately. ^36^

Specifically, the neck curvature (*M*_*ne*_) is determined as the local curvature at the contact point, as given below, namely *M*_*LS*_ between the mother (large) and interior daughter (small) vesicles, and *M*_*SS*_ between the daughter (small) - daughter (small) vesicles ^10^.

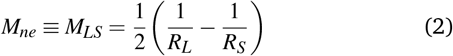

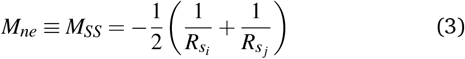

The symbols *R*_*L*_ denote the mother vesicle radius, *R*_*S*_ denotes the daughter vesicle radius, *R*_*S,i*_ and *R*_*S, j*_ are used for the radii of individual daughter buds at a given time, and *i* ≠ *j* ^10^. The negative sign of (−1*/R*_*s*_) is used for inbuds and a positive sign for (1*/R*_*L*_). The dynamic changes in these metrics reveal the temporal evolution of the local membrane curvature during vesicle growth.

Both curvature metrics remain negative throughout the recorded budding window, consistent with the inward-curving neck geometry captured by the sign convention in Eqns. (2)– (3). *M*_*SS*_ initially shows an increase in magnitude within roughly the first 20 − 30% of the normalised time, after which the curvature plateaus (1,1.5 µM) or, at higher concentrations, undergoes a further non-monotonic decrease before recovering in the later times (most prominently at 2 µM for both lipid compositions). Whereas *M*_*LS*_ typically begins close to zero at early *t/t*_max_ and decreases progressively as budding proceeds, indicating that the mother–bud neck continues to tighten well after the daughter compartment first becomes resolvable as a distinct object, rather than adopting a fixed geometry immediately upon formation. This distinction suggests that mother–bud necks (*M*_*LS*_) tighten quasi-monotonically over the course of budding, whereas bud– bud contacts (*M*_*SS*_) are established relatively rapidly and then continue to reorganize as neighboring buds compete for larger membrane area and contact geometry. *M*_*SS*_ magnitudes are also substantially larger than *M*_*LS*_ throughout (~ 0.1–0.9 *µ*m^−1^ versus ~ 0.02–0.35 *µ*m^−1^), as expected from Eqs. (2)–(3); which shows *M*_*SS*_ is calculated as the summation of two small-radius curvature contributions, whereas *M*_*LS*_ is the difference between the mother-vesicle curvature and daughter-vesicle curvature, that are comparatively smaller in magnitude. Comparing lipid compositions, POPC–Chol vesicles reach larger neck-curvature magnitudes than POPC at matched alpha-amylase concentration (e.g., *M*_*SS*_ approaching −0.9 *µ*m^−1^ for POPC–Chol versus ~ −0.7 *µ*m^−1^ for POPC at 2.5 *µ*M, consistent with cholesterol’s known condensing effect on the bilayer favouring tighter, smaller-radius neck geometries. Similarly, for the same lipid composition, an increase in alpha-amylase concentration increases the magnitude of both neck curvature metrics, *M*_*SS*_ and *M*_*LS*_, as observed from Figures 8 and 9. A colormap showing the variation in density of *M*_*SS*_ and *M*_*LS*_ with the normalized time duration across the four different alpha-amylase concentrations for both POPC and POPC-Chol compositions is shown in Supplementary Figures 7 and 8. The intensity of colour highlights the relative density of the curvature metrics, which provides us with useful information about how the specific magnitude of both *M*_*SS*_ and *M*_*LS*_ vary for different mother-daughter and daughter-daughter interactions within the same time-point for each vesicle.

**Fig. 8.**
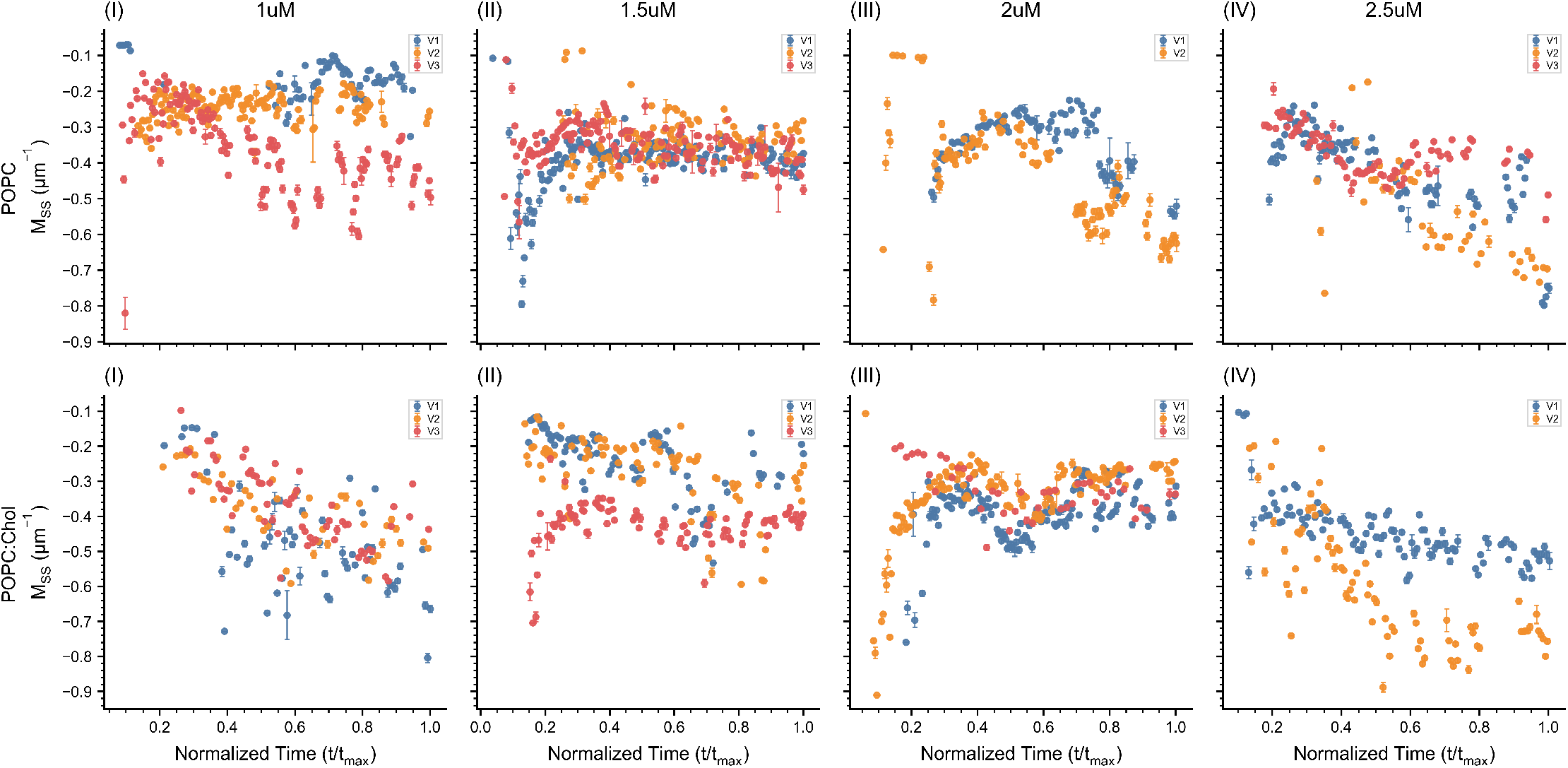
Temporal evolution of *M*_*SS*_ during vesicle budding. The neck curvature between pairs of in-bud daughter vesicles (*M*_*SS*_) is plotted as a function of normalized time (*t/t*_max_) for representative POPC (top-row) and POPC–Chol (bottom-row) vesicles. Subpanels (I)–(IV) correspond to alpha-amylase concentrations of 1, 1.5, 2, and 2.5 *µ*M, respectively. Data from individual vesicles (V1–V3) are shown as 2-minute binned mean values with error bars representing the standard deviation within each bin. Time was normalized by the total duration of the budding event (*t*_max_), allowing comparison of the temporal evolution of daughter–daughter neck curvature across lipid compositions and experimental conditions.

**Fig. 9.**
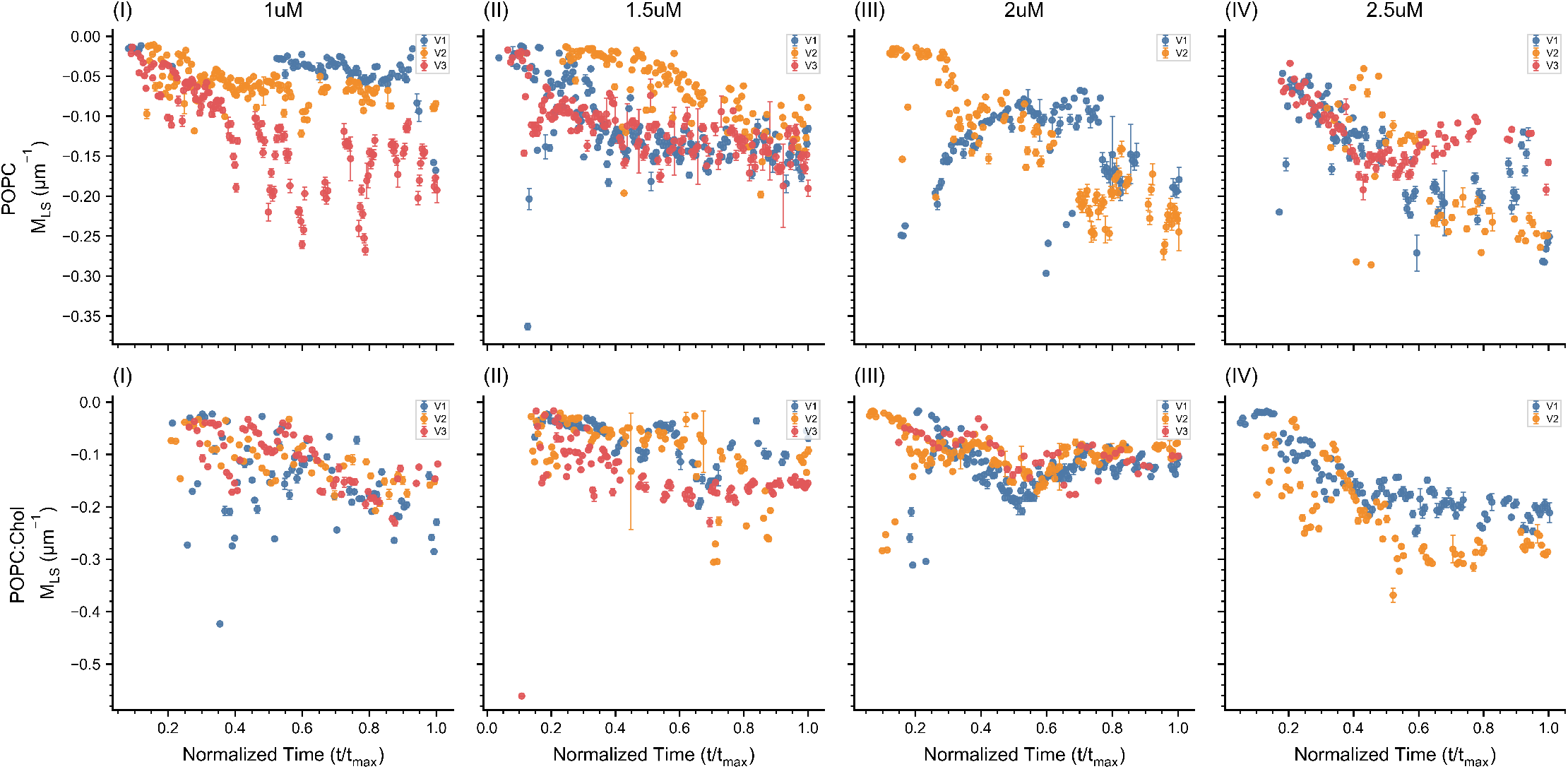
Temporal evolution of *M*_*LS*_ during vesicle budding. The neck curvature between the mother vesicle and its in-bud daughter vesicle (*M*_*LS*_) is plotted as a function of normalized time (*t/t*_max_) for representative POPC (top-row) and POPC–Chol (bottom row) vesicles. Subpanels (I)–(IV) correspond to alpha-amylase concentrations of 1, 1.5, 2, and 2.5 *µ*M, respectively. Data from individual vesicles (V1–V3) are shown as 2-minute binned mean values with error bars representing the standard deviation within each bin. Time was normalized by the total duration of the budding event (*t*_max_), enabling comparison of the temporal evolution of mother–daughter neck curvature across lipid compositions and experimental conditions.

While Figures 8 and 9 provide us with useful information about the evolution of neck curvature metrics and the interaction between mother and daughter vesicles during the time course of the experiment, they do not provide any spatial information about the specific locations about these interaction events. A spatio-temporal analysis precisely highlighting when and where these daughter–daughter or mother–daughter contacts occur within the vesicle can provide valuable information regarding the dynamics of such morphological changes.

To quantify these events spatio-temporally, we created polar plots to map the curvature metrics *M*_*SS*_ and *M*_*LS*_ within the detected mother-vesicle contour as shown in Figure 10. In this representation, the angular coordinate marks the azimuthal position of the contact point along the mother-vesicle boundary, while the radial coordinate corresponds to the mother-vesicle radius at the time that contact was recordedranging from the initial (outermost) radius to the final, most-deflated (innermost) radius, with concentric gridlines labelled in micrometers. For both sets of lipid compositions and the four different alpha amylase concentrations shown in Figure 10 a and b, the polar plots for each group show to distinct color-bars as highlighted by two different rows. The top row of each figure colour-codes the neck curvature magnitude (*M*_*SS*_ and *M*_*LS*_ respectively) at each contact point, and the bottom row colour-codes the recording time explicitly, denoting the time at which the contact between the mother-daughter and daughter-daughter vesicles are made. Since the mother vesicle deflates monotonically over the course of the experiment, we observe that the contacts near the outer edge of the vesicle occur primarily in the early phase of the experiment, at initial stages of budding, while with increase in time the contact points increasingly drift towards the center which highlights the reduction in mother vesicle area over time. Our results show that across the different concentrations and lipid compositions, contact points are broadly distributed around the full angular range of the mother-vesicle contour, with no consistent preferred angular position or persistent hotspot across conditions. At higher alpha-amylase concentrations (2.5 *µ*M), few individual vesicles show some local angular enrichment leading to asymmetries in the distribution of contact point locations; however these patterns are not consistently preserved across all the different concentrations or between the two lipid compositions, being POPC and POPC–Chol. Thus, the spatial maps provide no clear evidence for a strong contour-localized nucleation site or systematic angular templating of new buds (daughter vesicles) by previously formed buds. Instead, budding contacts appear to be distributed over the available mother contour, with local deviations from uniformity likely reflecting heterogeneity between individual budding events. The time-coded maps further show a broadly radial progression of contact formation, with earlier contacts occupying the outer regions and progressively later contacts appearing at smaller radii as the mother vesicle deflates. This correspondence is particularly apparent in several of the POPC conditions, where successive time populations form approximately concentric bands, although the later stages become more spatially heterogeneous in some vesicles. The *M*_*SS*_ maps are generally more spatially and angularly uniform than the *M*_*LS*_ maps, whereas *M*_*LS*_ shows more pronounced ring-like organization together with relatively local angular heterogeneity. Collectively, these results offer important insights into the dynamic coupling between mother vesicles and their developing in-bud daughters during shape evolution, capturing progressive changes in local membrane curvature during budding that are likely closely tied to local variations in membrane mechanics and to the alpha-amylase-driven osmotic deflation rate of the mother vesicle.

**Fig. 10.**
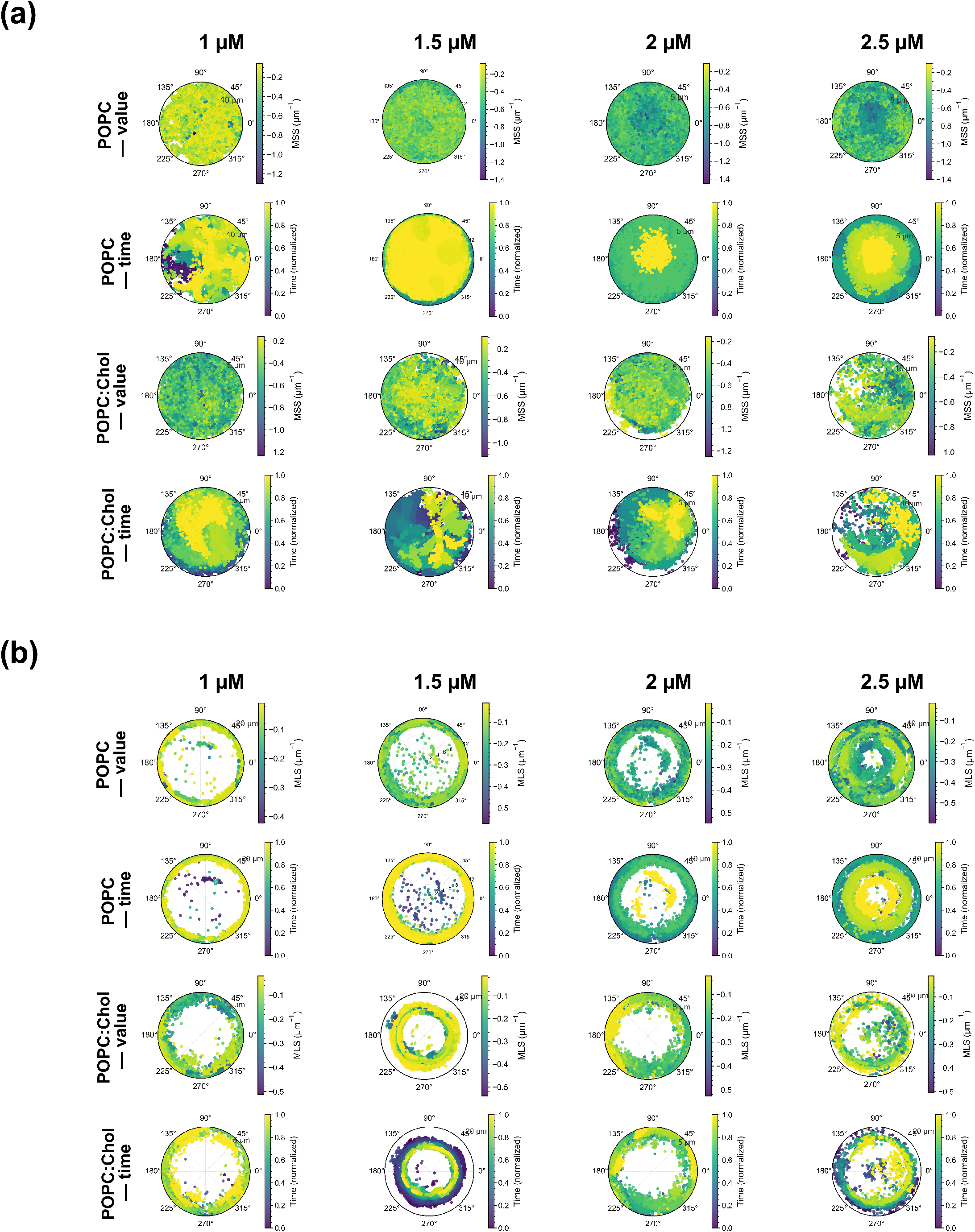
Polar maps showing the spatial and temporal distribution of contact curvature for POPC and POPC–Chol GUVs at 1–2.5 µM alpha-amylase. Columns correspond to increasing alpha-amylase concentration. In each map, points are plotted at their measured angular position *θ* and radial distance from the mother-vesicle centroid. The upper and lower sets of rows show curvature magnitude and acquisition time, respectively, thereby resolving both the spatial distribution and temporal evolution of contact regions during vesicle remodeling. (a) Daughter–daughter contacts quantified using the local-secant curvature, *M*_*SS*_. (b) Mother–daughter contacts quantified using the local-secant curvature, *M*_*LS*_.

### 4.4 Bimodal distribution curvature populations

To further quantify the pooled distributions of the neck-curvature metrics, *M*_*SS*_ and *M*_*LS*_ across the lifespan of each vesicle, we fitted a two-component Gaussian mixture model to each datasets (Supplementary Figures 9 and 10). For each vesicle, the binned curvature values were pooled over the recording and represented as normalized histograms with kernel-density estimates, and the resulting distributions were fitted with a two-component Gaussian mixture model. Our experimental results suggest a bimodal distribution for both curvature metrics as they can be resolved into two statistically distinct curvature populations across essentially all tested concentrations and lipid compositions. The corresponding fitted population means, standard deviations, and mixture weights (*w*) for each vesicle datasets are summarized in Supplementary Table 2. To construct these distributions, we used measurements that were binned every 2 minutes and hence they can capture the entire range and relative occurrence of curvature values sampled during the recording rather than individual events.

The persistent presence of a bimodal curvature distribution across the entire tested population range indicates that vesicle neck regions occupy two discrete geometric regimes during remodeling; with plausibly a population of actively growing necks (higher-magnitude curvature) coexisting with necks that have relaxed toward a more mechanically stable radius (lower-magnitude curvature), rather than continuously sampling a single distribution of neck geometries. Although the mean curvature values of *M*_*SS*_ and *M*_*LS*_ differ in magnitude, their overall distributions display a comparable bimodal pattern across the different concentrations and lipid compositions (Supplementary Figure 9– 10). However, the relative weighting of the two populations differs for individual samples within each subgroups and do not occur with equal frequencies. The fitted mixture weights (*w*) in Supplementary Table 2 suggests that for each vesicles one curvature population tends to systematically dominate under particular alpha-amylase concentrations or lipid compositions.

For both *M*_*SS*_ and *M*_*LS*_, the fitted components generally correspond to a higher-magnitude and a lower-magnitude curvature population, although the degree of separation varies substantially between conditions and individual vesicles. The distinction is most apparent for *M*_*SS*_, particularly at higher alpha-amylase concentrations, where the two fitted means become increasingly separated in several vesicles. In POPC, for example, the highermagnitude population shifts from approximately −0.18 *µ*m^−1^ at 1 µM to values as large in magnitude as −0.77 *µ*m^−1^ at 2.5 µM, while the second population remains at comparatively lower magnitudes. POPC–Chol shows a similar tendency, with the higher-magnitude population reaching approximately −0.67 to −0.71 *µ*m^−1^ in several vesicles at the higher concentrations. The histograms likewise show that the two populations are not equally well resolved in every condition: some distributions contain strongly overlapping peaks, whereas others exhibit a much clearer separation between the lower- and higher-curvature regimes.

The relative weights of the two populations also vary considerably between vesicles and experimental conditions, rather than following a single concentration-dependent trend. In POPC, several vesicles at 1 and 1.5 µM show relatively balanced contributions from the two components, whereas at 2.5 µM some vesicles are strongly dominated by the lower-magnitude population. At the same time, V1 at 2.5 µM contains a small but clearly separated higher-magnitude *M*_*SS*_ component, with a fitted weight of only *w*_1_ = 0.054. POPC–Chol shows similarly heterogeneous weighting: the lower-magnitude population dominates several vesicles at 1.5 and 2 µM, while the higher-magnitude population becomes dominant in the 2.5 µM V1 vesicle, where it accounts for approximately 92% of the fitted *M*_*SS*_ distribution. Thus, although two curvature ranges recur across the dataset, their relative representation is strongly vesicle- and conditiondependent. *M*_*LS*_ shows the same general two-population structure but over a substantially smaller curvature range than *M*_*SS*_. The higher-magnitude population generally lies between approximately −0.07 and −0.28 *µ*m^−1^, whereas the lower-magnitude population is typically closer to zero, often between approximately −0.02 and −0.12 *µ*m^−1^. The histogram distributions show that these populations are sometimes strongly overlapping, particularly at lower concentrations, while clearer separation becomes apparent in several of the higher-concentration conditions. The mixture weights again vary between vesicles, with some distributions dominated by the lower-magnitude component and others showing comparable contributions from both components. POPC–Chol displays particularly strong vesicle-to-vesicle variation, with no consistent monotonic shift in the relative population weights across alpha-amylase concentration.

Taken together, the 2-min binned distributions reveal recurring low- and high-magnitude curvature ranges during budding, with the separation between these ranges generally more pronounced for *M*_*SS*_ than for *M*_*LS*_.The concentration- and lipid-dependent changes of the two population means in both *M*_*SS*_ and *M*_*LS*_ is examined quantitatively in Figure 11. Our results show a gradual increase in the magnitudes of the curvature means with increase in concentrations which can be attributed to higher membrane leaflet asymmetry. The shift of the higher-magnitude *M*_*SS*_ population toward stronger curvature at several higher alpha-amylase concentrations is consistent with the development of tighter bud– bud neck geometries under stronger osmotic deflation. However, the variation in mixture weights and the substantial overlap between fitted components in several conditions indicate that these should not be interpreted as strictly discrete or permanently occupied curvature states. Rather, the Gaussian-mixture analysis provides a compact description of the heterogeneous curvature values sampled over the course of budding. In this sense, the analysis complements the temporal and spatial measurements by showing that the evolving neck geometry spans distinct but overlapping curvature regimes, whose relative representation changes between vesicles, lipid compositions, and alpha-amylase concentrations.

**Fig. 11.**
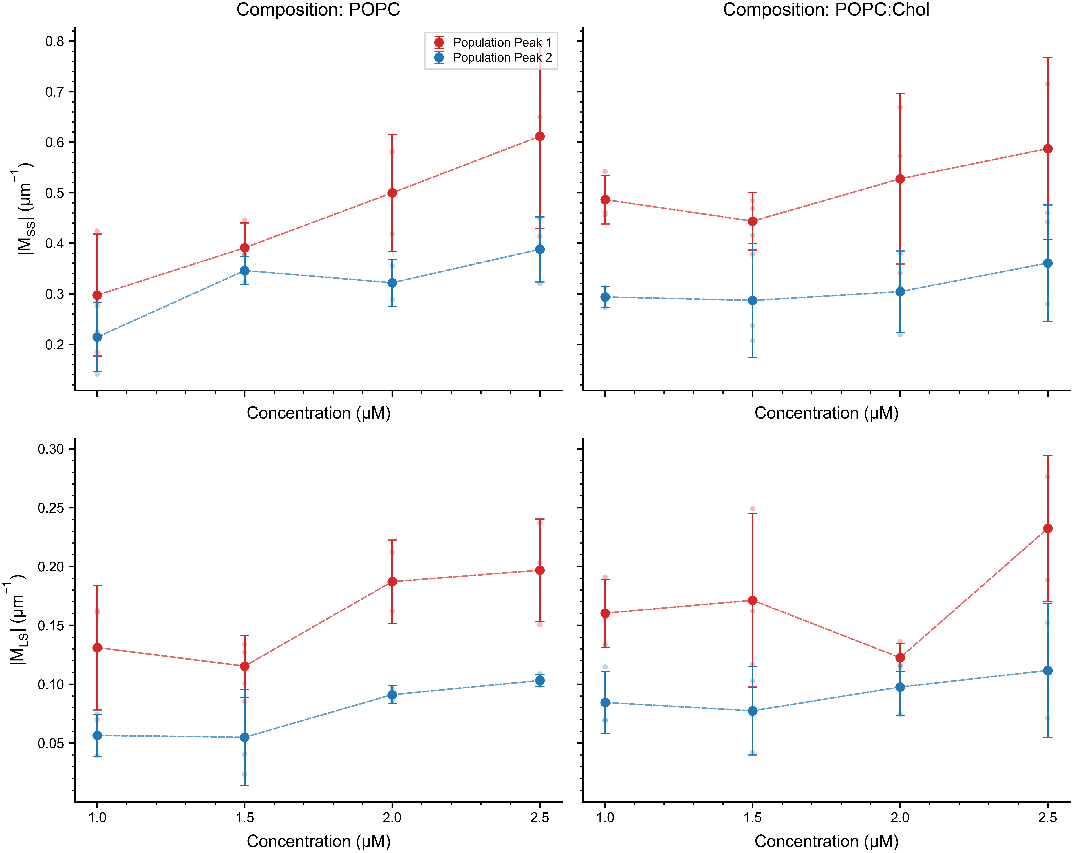
Concentration- and lipid-dependent variation of the two fitted curvature populations for *M*_*SS*_ (top) and *M*_*LS*_ (bottom). Points represent the mean curvature magnitude of Population 1 and Population 2 obtained from the two-component Gaussian-mixture fits, with error bars indicating the corresponding variability across vesicles. Results are shown separately for POPC and POPC–Chol GUVs across 1–2.5 µM alpha-amylase.

### 4.5 Time scale of Mother-vesicle shape evolution are concentration-dependent

The present work focuses on strongly driven enzyme-mediated osmotic remodeling. To quantify the temporal dynamics of mothervesicle deflation, under weak perturbations, where intrinsic shape relaxation could be separated from external chemical driving. We fitted the normalized mother-vesicle area, *A*(*t*)*/A*_0_, with an exponential decay model of the form :-

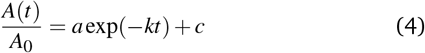

where *k* represents the characteristic decay rate, and the corresponding characteristic time scales is given by *τ* = 1*/k*. The mother-vesicle area exhibit a well-defined relaxation behavior, during the transition from a quasi-spherical vesicle to a multibudded state under osmotic stress. The fitted trajectories values are shown in Figure 12, with the resulting parameters summarized in Supplementary Table 3. The process is externally driven by enzymatic hydrolysis in the outer medium.

**Fig. 12.**
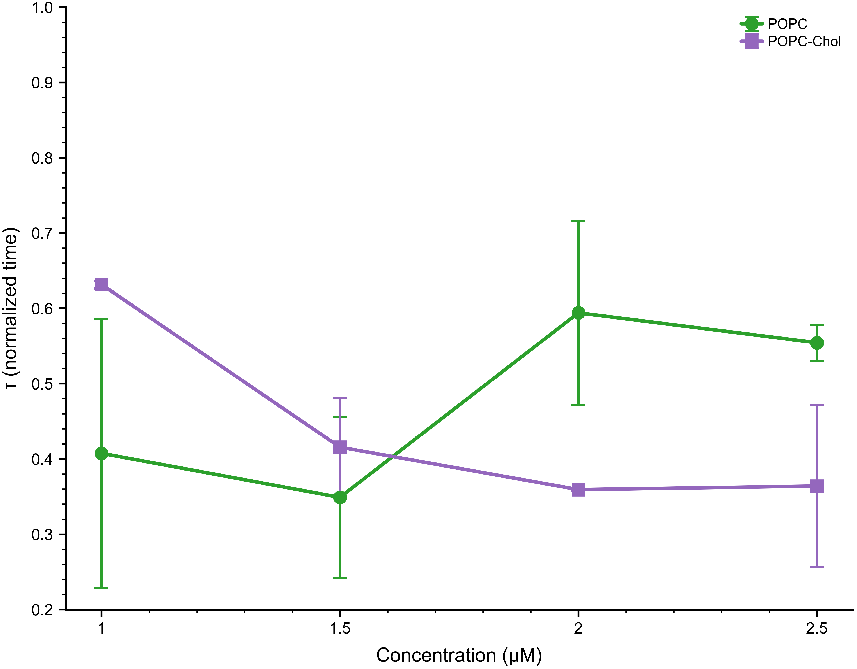
Exponential fit to the normalized mother-vesicle area decay as a function of normalized time for POPC and POPC–Chol GUVs at 1– 2.5 µM alpha-amylase. The characteristic time scale, *τ* obtained from the exponential fits are shown for each lipid composition and alpha-amylase concentration. Symbols and error bars represent the mean and standard deviation across vesicles, respectively, with solid and dashed lines indicating the corresponding trends for the POPC and POPC-Chol vesicles.

The characteristic time scale shows a non-monotonic relationship with alpha-amylase concentration for both lipid types. For POPC, *τ* is about 0.41 at 1 *µ*M (n = 2), 0.35 at 1.5 *µ*M (n = 3), 0.59 at 2 *µ*M (n = 2) and 0.55 at 2.5 *µ*M (n = 3). Given the vesicle-to-vesicle spread (Supplementary Table 3), the values at 1 and 1.5 *µ*M are indistinguishable, and there is at most a modest increase in *τ* at 2–2.5 *µ*M. For POPC–Chol, *τ* decreases from about 0.63 at 1 *µ*M to 0.42 at 1.5 *µ*M and 0.36 at 2 *µ*M, and then remains at about 0.36 at 2.5 *µ*M (n = 2, with larger uncertainty). Thus, POPC–Chol shows a decreasing trend of *τ* with alpha-amylase concentration that levels off at higher concentrations, whereas POPC shows no clear monotonic trend within the experimental uncertainty. The time scales of the two compositions overlap at 1.5 *µ*M and differ most at 2–2.5 *µ*M, where POPC–Chol has the smaller *τ*. Because *τ* is expressed in time normalized by *t*_max_ and is based on only two to three vesicles per condition, these differences should be regarded as indicative rather than conclusive; they suggest, but do not by themselves establish, that the deflation kinetics depend on both alpha-amylase concentration and lipid composition. The single-exponential model captures the main relaxation time of mother-vesicle area loss, but it does not identify a specific physical mechanism. Its advantage is that it describes the normalized area curves with few parameters and allows a simple comparison of deflation dynamics across lipid compositions and alpha-amylase concentrations. Previous work has described membrane processes in terms of relaxation and cooperative dynamics, including studies connecting lipid transfer kinetics to characteristic relaxation times derived from thermodynamic parameters ^37^.

## 5 Discussion

In this study, we present a customized automated image-analysis workflow, guided by physical constraints, to quantify budding dynamics in individual giant unilamellar vesicles using phase-contrast time-lapse imaging. This approach integrates contourbased detection of the mother vesicle with machine-learningbased instance segmentation of daughter compartments, allowing accurate identification and tracking of individual buds at high temporal resolution over extended periods, even in low-contrast images and under evolving membrane shapes.

As illustrated in Figure 1, the tailored image-segmentation pipeline robustly tracks and quantifies vesicle shape changes throughout the experiment and across multiple independent data sets. To further highlight the robustness of our framework in accommodating diverse initial mother–daughter (in-bud) vesicle morphologies, we analyzed multiple recordings obtained from vesicles with distinct initial geometries and at different time points. Moreover, tracking individual daughter compartments reveals a variety of dynamical behaviors, including transient buds, long-lived daughter vesicles, and short-lived contact events with the mother membrane or between neighboring daughters. By combining geometry-based filtering with contour-level contact detection, we can reliably distinguish which of these events correspond to physically meaningful membrane remodeling and filter out artifacts arising from segmentation errors. Within a subset of frames, segmentation inaccuracies in daughter regions produced by pre-trained models suggest that a dedicated CNN-based segmentation strategy, such as StarDist or Cellpose, could more faithfully capture these features which further reinforces our choice to incorporate these tools within our custom framework. We also validated the model performance by artificially inducing noise into the system and observed that the segmentation pipeline could provide reliable results up to 15% noise highlighting the robustness of the chosen framework as shown in Figure 6

The daughter-vesicle segmentation was performed using pre-trained StarDist and Cellpose-SAM models, without additional training or fine-tuning on GUV-specific images. The criteria used for mother-vesicle detection are also described in greater detail, including the scoring based on circularity, vesicle size, and temporal consistency. We have described the optional Richardson–Lucy deconvolution step and the use of a Gaussian point-spread function. For quantitative validation, we now report IoU, daughtervesicle precision, mother-vesicle area agreement, centroid error, Hausdorff discrepancy, and vesicle-quantification deviation. We include a systematic noise-robustness analysis to characterize the performance of the pipeline under different image-quality conditions. Finally, to improve reproducibility, we have made the complete analysis code publicly available through Zenodo.

In addition to identifying vesicle contours as depicted in Figure 1, segmented images were utilized to quantify changes in the equatorial area shown in Supplementary Figure 3 within individual time-lapse datasets.With time, a significant decrease in the mother vesicle size was observed, suggesting that enhanced enzymatic hydrolysis accelerated osmotic deflation of the mother vesicle. The rate of starch hydrolysis by AA (outside the GUVs) determines the magnitude of sugar accumulation in the bulk solution. Reducing sugar concentration increased progressively over time, as shown in Figure 4. The temporal progression of *A* for both mother vesicle and inbuds across three individual GUVs is shown in Figure 7. The dynamics of shape evolution of the mother vesicle area was captured using a simple exponential relaxation model (Eqn. 4); which reveals the existence of a characteristic time scale that dictates the vesicle relaxation towards the final contracted shape. The broadly overlapping time scales observed for POPC and POPC–Chol suggest that lipid composition does not produce a uniform shift in deflation kinetics as observed in Figure 12.

Analyzing the mean curvature of mother and daughter vesicles shown in Figures. 9 and 8 offers valuable insights into their local bending over time, which are essential to understand the physical mechanisms driving these responses. Measurement of local curvature at contact points between mother and in-bud vesicles provides key information on changes in membrane geometry at these sites, thus connecting observed morphological alterations to the mechanical properties of the membrane that cause them. Further quantification of the local geometry changes during mother– daughter and daughter–daughter contacts were performed using the neck curvature metrics *M*_*LS*_ and *M*_*SS*_, respectively. Both metrics remained predominantly negative, consistent with the inward-curving geometry of the budding necks. *M*_*LS*_ generally became progressively more negative during budding, whereas *M*_*SS*_ showed a more rapid initial increase in magnitude followed by plateaus or non-monotonic changes, indicating that mother– daughter necks tighten progressively while daughter–daughter contacts undergo further geometric reorganization during budding. *M*_*SS*_ also reached substantially larger magnitudes than *M*_*LS*_, reflecting the different geometric contributions entering the two curvature definitions. At matched alpha-amylase concentrations, POPC–Chol vesicles reached larger |*M*_*SS*_| values than POPC vesicles in several conditions, indicating that lipid composition influences the local neck geometries attained during budding.

Mapping these curvature measurements onto the mother-vesicle contour further showed that contact points were broadly distributed over the available angular range, with no consistent preferred angular position or persistent hotspot across concentrations and lipid compositions. Although individual vesicles displayed local angular enrichments, these patterns were not consistently preserved across conditions, providing no clear evidence for systematic angular templating or a strongly localized nucleation site. Instead, the spatial maps showed a broadly radial progression of contacts as the mother vesicle deflated, with earlier contacts occurring at larger radii and later contacts appearing progressively closer to the contracted centre. Additional analysis of the 2-minute binned curvature distributions revealed a bimodal distribution for both *M*_*SS*_ and *M*_*LS*_ across the tested concentrations and lipid compositions. The higher-magnitude population became more negative for *M*_*SS*_ under several of the higher alpha-amylase conditions, while the relative weights of the two populations varied between individual vesicles and did not follow a single universal concentration-dependent trend. These results indicate that the evolving neck geometry samples distinct lower- and higher-curvature regimes during budding, with their relative occurrence changing between vesicles and experimental conditions.

The potential effect of manual region of interest selection on reproducibility is discussed. The purpose of the ROI selection in our framework is to restrict the analysis to the target GUV, exclude any neighboring structures and therefore reduce the size of the image array used for subsequent analysis. The ROI can also be adjusted during long acquisitions to accommodate for vesicle drift. Currently the ROI acts as an initial spatial constraint; which influences the subsequent mother-vesicle detection, daughter-vesicle segmentation, object filtering and temporal tracking. To quantify the sensitivity of this step, we repeated the ROI selection 10 times on the same representative frame, using separate, independently selected ROI boundaries, and evaluated the resulting variation in the downstream segmentation. The resulting mother-area deviations are shown in Supplementary Figure 2e. The repeated measurements were clustered within a relatively narrow range, with the measurements centered at approximately 5% mother-area de viation, providing an estimate of the variability introduced by re peated ROI selection under the tested conditions. This analysi characterizes the sensitivity of the workflow to the manual RO boundary rather than claim that the ROI-selection step is com pletely operator-independent.

We have now clarified the scope of the automation in the revised manuscript. The current implementation is not claimed to be fully automated from raw image acquisition onward, because the ini tial ROI selection remains interactive. Instead, the ROI provides a user-defined spatial constraint, after which the mother-vesicle detection, daughter-vesicle segmentation, object filtering and temporal tracking are performed computationally. We acknowledge that the removal of this initial manual step would further improve scalability and have now described the ROI selection explicitly as an interactive initialization step rather than representing the entire workflow as fully automated in the revised manuscript.

Overall, this study demonstrates a practical and reproducible strategy for extracting quantitative, time-resolved geometric in formation from short and long vesicle movies, where ensemble averaging is not possible, and physical consistency must be enforced at the level of a single object. Taken together, these results connect the biochemical activity of alpha-amylase in the externa medium to the progressive osmotic deflation of the mother vesicle and the accompanying remodeling of daughter compartments. The combined analysis of global area dynamics, deflation kinetics, time-resolved neck curvature, spatial contact mapping, and curvature distributions shows that mother-vesicle contraction and daughter-vesicle remodeling are dynamically coupled processes, while also revealing substantial heterogeneity between individual vesicles and experimental conditions. The proposed framework is readily adaptable to static images or long-term time-series data, different lipid compositions, or externally perturbed systems, and provides a useful bridge between time-resolved imaging experiments and theoretical descriptions of curvature-driven membrane remodeling in soft-matter systems.

The extracted parameters, such as vesicle area and local curvature, are not connected to a mechanistic interpretation based on established frameworks (e.g., membrane elasticity theory, Helfrich energy, ADE theory), in this paper. The study is relevant to the Soft Matter readership, and the development of automated tools for analyzing complex vesicle morphologies is valuable. The methodology appears promising, particularly for handling crowded and evolving structures in phase-contrast microscopy. The study presents an automated image-analysis method to quantify shape transformations in giant unilamellar vesicles (GUVs), combining contour-based detection with deep learning segmentation and tracking. The topic could be of potential interest for the readership of Soft Matter and, more broadly, for scientists studying dynamical changes in GUVs subjected to environmental stimuli. The integration of dynamical detection with machinelearning tools, in principle, could provide a useful bridge between experiments and theoretical descriptions of membrane remodeling.

## Author contributions

T.P. contributed to code design, data analysis, figure preparation, and paper writing. R.K. conducted experiments and prepared experimental figures. J.V.B.S. handled coding and data analysis for curvature mapping and prepared SI Figure 4. C.D. conducted experiments for the images shown in SI Figure 4. S.N and D.S. contributed to the HPLC experiment and shared Figure 3 with the text. A.C. contributed to the code design, analysis, interpretation, and manuscript writing. T.B. participated in designing the experiments, collecting and analyzing the data, interpreting the results, and preparing the manuscript. All authors reviewed and approved the final manuscript.

## Data availability

All relevant data are included within the manuscript. Any additional information is available from the authors upon reasonable request. The code is available at Zenodo: https://zenodo.org/records/22210498.

## Acknowledgements

This research was conducted within the Indian Institute of Science Education and Research Mohali and supported by the Ramalingaswami grant of DBT (DBT/RLF/Re-entry/06/2020), DST/INT/DAAD/P-1 6/2023 (G) and IC-12025(22)/1/2023-ICD-DBT. Research Associate fellowship for C.D. is supported by DBT/RLF/Re-entry/06/2020 grant money. AC acknowledges the_1_ intramural financial support from IIT Guwahati.

